# Oxford Nanopore and Bionano Genomics technologies evaluation for plant structural variation detection

**DOI:** 10.1101/2021.04.16.440130

**Authors:** Aurélie Canaguier, Romane Guilbaud, Erwan Denis, Ghislaine Magdelenat, Caroline Belser, Benjamin Istace, Corinne Cruaud, Patrick Wincker, Marie-Christine Le Paslier, Patricia Faivre-Rampant, Valérie Barbe

**Author notes:** Corresponding author: Patricia Faivre Rampant.

## Abstract

**Background:** Structural Variations (SVs) are very diverse genomic rearrangements. In the past, their detection was restricted to cytological approaches, then to NGS read size and partitionned assemblies. Due to the current capabilities of technologies such as long read sequencing and optical mapping, larger SVs detection are becoming more and more accessible.

This study proposes a comparison in SVs detection and characterization from long-read sequencing obtained with the MinION device developed by Oxford Nanopore Technologies and from optical mapping produced by the Saphyr device commercialized by Bionano Genomics. The genomes of the two *Arabidopsis thaliana* ecotypes Columbia-0 (Col-0) and Landsberg *erecta* 1 (L*er*-1) were chosen to guide the use of one or the other technology.

**Results:** We described the SVs detected from the alignment of the best ONT assembly and DLE-1 optical maps of *A. thaliana* L*er*-1 on the public reference Col-0 TAIR10.1. After filtering, 1 184 and 591 L*er*-1 SVs were retained from ONT and BioNano technologies respectively. A total of 948 L*er*-1 ONT SVs (80.1%) corresponded to 563 Bionano SVs (95.3%) leading to 563 common locations in both technologies. The specific locations were scrutinized to assess improvement in SV detection by either technology. The ONT SVs were mostly detected near TE and gene features, and resistance genes seemed particularly impacted.

**Conclusions:** Structural variations linked to ONT sequencing error were removed and false positives limited, with high quality Bionano SVs being conserved. When compared with the Col-0 TAIR10.1 reference, most of detected SVs were found in same locations. ONT assembly sequence leads to more specific SVs than Bionano one, the later being more efficient to characterize large SVs. Even if both technologies are obvious complementary approaches, ONT data appears to be more adapted to large scale populations study, while Bionano performs better in improving assembly and describing specificity of a genome compared to a reference.

## Background

Structural variations (SV) are defined as genomic variations involving segments of DNA from 50 bp to several megabases. SVs consist of unbalanced rearrangements such as copy number variations (CNV) including insertions/deletions (Indels) and presence/absence variations (PAV), and balanced events like inversions and translocations [1,2,3,4]. Several mechanisms explain the formation of SVs, such as recombination errors generated by non-homologous end-joining and non-allelic homologous recombination, genome duplication and transposition [1,2]. The structural variations in human were largely studied and recently, Ho *et al*. reviewed the impact of the SVs in human deseases [4]. In plants it has been shown that the structural variations play a key role in evolution of genomes and are responsible for phenotypic variations by impacting TEs and genes [3,5,6,7,8]. In particular, SVs were found in stress related and resistance genes [9,10,11,12,13], to be related to local adaptation [14,15], or linked to other traits of agronomical interest such as tomato fruit flavor, rice grain size or poplar wood formation [16,17,18].

Nowaday, identification of SVs contributes to the construction of the Panreference genome or super pangenome [19,20]. This new approach to build a reference will better reflect the genetic diversity of the species, and in the same time deepen the understanding of genome evolution, as well as enhancing knowledge of adaptative traits [21,22,23,24,25].

The development of new sequencing technologies has boosted studies of SVs found in a genome, which were detected until recently only by CGH arrays or SNP [26,27,28]. Short read sequencing technologies have made possible the identification of SVs in several species [29,30,31,32,33,34,35,36]. However, the size of the reads is a limiting factor for the detection of large SVs and SVs in highly repetitive regions. The 3^rd^ generation sequencing offer new opportunities to identify SVs at larger scale with two kinds of methods. First are based on linked short reads, as in 10x Genomics and Hi-C approaches [37], second by directly generating long reads, as proposed by Pacific Biosciences [38] and Oxford Nanopore Technologies (ONT) [39,40]. These approaches provide access to complex regions, increasing their uses to produce genome assemblies and to detect structural variations in human [4,41,42,43], in *Arabidopsis thaliana* [24,44,45] and in other plants [46,47]. In parallel, a technology based on physical map and developed by Bionano Genomics [48], generates information from very large DNA molecules. These maps, named optical maps, are frequently generated to improve and validate sequencing assembly, to detect SVs in human genomes [43,49,50,51,52] and more recently in plants [7,46,47,53]. These 3^rd^ generation sequencing data made possible the identification of genetic rearrangements between individuals at intra specific level [53].

Herein, we obtained draft assemblies using Oxford Nanopore technology and Bionano Genomics optical maps, in order to compare the detection and characterization of the structural variations by both methods. Despite comparisons between two sequencing technologies or SV detection softwares are not anymore an uncharted territory [24,43,44,54], the comparison of two fundamentaly different technologies like ONT and Bionano was only performed in animals (Chimpanze [52] and Drosophila [55]), but not yet in plants. *A. thaliana* is a model organism with a small genome (130 Mb). For this study, we selected Columbia 0 (Col-0) and Landsberg *erecta* 1 (L*er*-1), two of the most studied ecotypes.

## Results

### ONT sequencing and genome assembly

The ONT sequence data of *Arabidopsis thaliana* ecotypes, Columbia (Col-0) and Landsberg *erecta* 1 (L*er*-1), were cleaned using the correction and trimming steps of Canu assembler [56]. A total of 9.8 Gb (N50=12.7 kb, 75X coverage) and 6.1 Gb (N50=16.5 kb, 47X coverage) were obtained for Col-0 and L*er*-1, respectively (Additional file 1: Tables S1 and S2).

To estimate ONT data completeness, the cleaned Ler-1 ONT reads were aligned against the L*er*-1 reference sequence with Minimap2 [57]. A total of 98.9% of the L*er*-1 reference sequence was covered by ONT reads. These L*er*-1 ONT data were also mapped against the Col-0 TAIR10.1 genome that was 95.2% covered (Additional file 1: Table S3). Samtools *depth* tool was then used on the L*er*-1 ONT reads mapping against the Col-0 TAIR10.1 reference to estimate the coverage at each position. The average coverage of 100 kb windows was 46.9X, with depth fluctuations in centromeric regions (Fig. 1).

**Figure 1.**
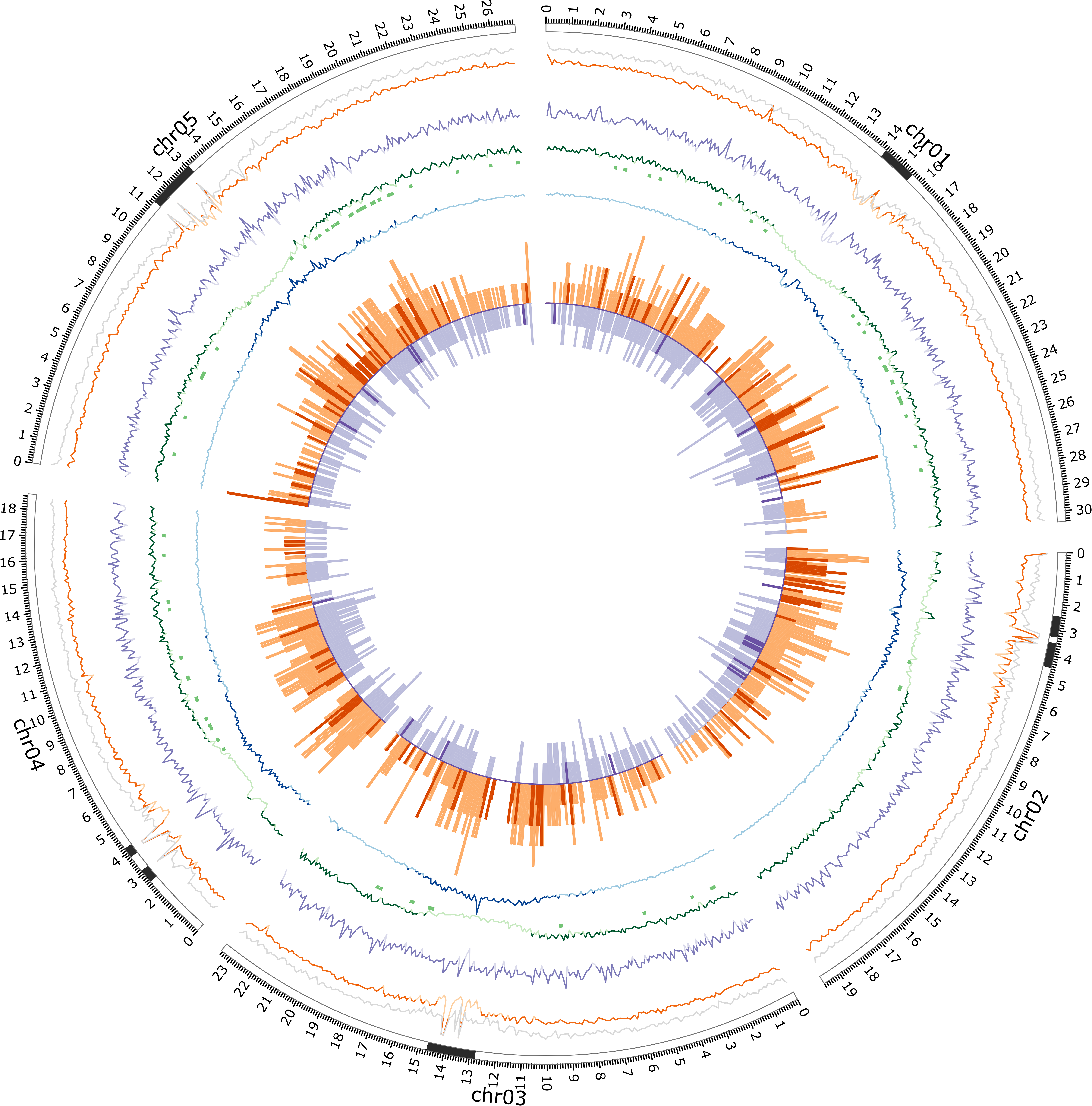
Circos visualization of L*er*-1 SVs landscape. All comparisons were performed against the Col-0 TAIR10.1 reference sequence per 100kb bins. From external to internal layer : Col-0 TAIR10.1 chromosomes (ticks every 100 kb) : black and light grey rectangles represent centromeric and NOR regions respectively; Average mapping coverage for Col-0 ONT reads (grey line) and L*er*-1 ONT reads (orange line with dark orange if coverage > 46X); DLE-1 label density as purple line (dark purple if density > 18 label per 100 kb); Genes density as green line (dark green if density > 23), NLR Genes [68] indicated as green rectangles; TEs density as blue line (dark blue if density > 58); ONT SVs occurences as orange outwards bars (dark orange bars represent ONT-specific SVs); Bionano SVs occurrences as purple inwards bars (dark purple bars represent Bionano-specific SVs).

To identify the best assembler for our data, *de novo* assemblies for Col-0 and L*er*-1 were performed with Canu, RA and SMARTdenovo (SDN). Based on general statistics (assembly size, contig number, N50 size), SMARTdenovo software generated better assemblies for both ecotypes compared to Canu or RA. (Additional file 1: Tables S4 and S5). Indeed, the SDN assemblies resulted in 79 contigs for Col-0 (cumulative size =117 Mb, N50=12.5 Mb with 5 contigs) and 101 contigs for L*er*-1 (cumulative sizes = 117 Mb, N50=10.7 Mb with 5 contigs). In addition, chimeric contigs were observed with Canu, while assemblies were more fragmented using RA (Additional file 2: Figures S1A-C and S2A-C). For all assemblers, centromeric regions were covered by many small contigs. These results were also supported by the alignements of the Col-0 and L*er*-1 assemblies on the respective reference chromosomes Col-0 TAIR10.1 [58] and L*er* [44]. The SDN assemblies named ONT Evry.Col-0 and Evry.L*er*-1 assemblies, were used to carry out subsequent SV analyses.

### Optical maps generation

The labelling of the genomic DNA was carried out using staining protocol with DLE-1 enzyme according to the manufacturer’s protocol. One run per ecotype on the Saphyr device was performed leading to 577.5 Gb and 610.9 Gb of molecules for Col-0 and L*er*-1 respectively. Molecules larger than 150 kb were selected leading to about 600-fold final coverage based on the theorical 130 Mb *Arabidopsis* genome size (Additional file 1: Tables S6 and S7). A total of 17 and 14 optical maps with N50 of 14.6 Mb and 14.7 Mb are generated for Col-0 and L*er*-1 respectively, bringing to a genome size of 125 Mb for both ecotypes (Additional file 1: Tables S8 and S9).

The average label density of the L*er*-1 optical maps was estimated at 18.47 per 100 kb (Additional file 1: Table S7). However, this DLE-1 density decreases in the centromeric regions due to molecule depth diminution and optical map breaks (Additional file 2: Figures S3A-E, Fig. 1).

### Structural Variations detection

The detections of the structural variations were performed independently using the ONT and Bionano technologies data and were carried out in two ways: L*er*-1 versus Col-0 TAIR10.1 reference and Col-0 versus L*er* reference. The different types of structural variations detected in our study are described in Additional file 2: Figure S4. Because general SVs characteristics (number, types and location) are similar in both types of analyses, only SV detection results from the Evry.L*er*-1 assembly and optical maps against the Col-0 TAIR10.1 reference are presented in details. Description of SVs detected by comparing the SDN assembly and optical maps Col-0 with L*er* reference are provided in Additional file 1: Tables S10-S14 and Additional file 2: Figures S5A-E.

The comparison of Evry.L*er*-1 assembly to Col-0 TAIR10.1 reference using MUMmer *show-diff* utility [59] revealed 2 186 potential SVs (Table 1).

**Table 1.** Types of Evry.L*er*-1 ONT and Bionano SVs obtained after alignement against Col-0 TAIR10.1 reference.

| SV Type | SEQ | BRK | JMP | TRA | INS/DEL/INV | TOTAL SV |
| --- | --- | --- | --- | --- | --- | --- |
| ONT | 108 | 6 | 5 | NA | 2 067 | 2 186 |
| Bionano | NA | NA | NA | 2 | 795 | 797 |

Note: NA : Not Available. SVs types are described in the Additional file 2: Figure S4.

A total of 119 SVs, called reference sequence junction (SEQ), break (BRK) and jump (JMP), found in centromeric, telomeric and nearby rDNA cluster were considered to correspond to unresolved assembly regions into Evry.L*er*-1 assembly compared to Col-0 TAIR10.1 reference and were filtered out.

To avoid false positive SV detection due to the ONT high error sequencing rate of 7.5% [60], a filter on query ONT structural variations size (> 1 kb) was applied. Out of the 2 186 SVs initially detected, 1 184 SVs remained (54.2% of total SVs) corresponding to 591 insertions (INS), 581 deletions (DEL) and 12 inversions (INV). No duplication was detected (Table 2).

**Table 2.** Characteristics of Evry.L*er*-1 ONT and Bionano SVs, obtained after alignement against Col-0 TAIR10.1 reference.

| Technology | ONT |  |  |  | Bionano |  |  |  |  |
| --- | --- | --- | --- | --- | --- | --- | --- | --- | --- |
| SV type | INS | DEL | INV | TOTAL | INS | DEL | INV | TRA | TOTAL |
| SV > 1kb (%) | 591 (49.9) | 581 (49.1) | 12 (1.0) | 1 184 | 289 (48.9) | 295 (49.9) | 5 (0.8) | 2 (0.4) | 591 |
| Cumulated Size | 3.4 | 4.0 | 0.3 | 7.7 | 2.9 | 2.3 | 1.6 | 0.4 | 7.2 |
| Median Size | 3 358 | 3 446 | 17 012 | 3 455 | 4 383 | 4 296 | 166 007 | 189 454 | 4 383 |
| Average Size | 5 724 | 6 801 | 26 865 | 6 467 | 10 021 | 7 885 | 310 470 | 189 454 | 12 104 |

Note : Cumulated sizes are in Mb, Median and average sizes in bp.

A 5 Mb insertion in the Evry.L*er*-1 assembly was detected on Chr3 Col-0 TAIR10.1 reference (14 272 986..14 284 724) due to a detection error of MUMmer in a complex region associated to a rDNA cluster. Thereby, this insertion was removed from the final data and not considered in the result. The ONT structural variations median size was 3 455 bp and the cumulated sizes 7.7 Mb. The SVs were equally distributed in size and number between INS and DEL. The INV categories had larger median and average sizes than INS and DEL. With a cumulated size of 0.3 Mb, INV represented 3.9% of the ONT variation size (Table 2). Structural variations were detected on all chromosomes, with a preferential location on chromosome arms and with no confident SV on the Chr1, 3 and 4 centromeres (Fig. 1).

Bionano data analysis (optical maps construction and SVs detection) was carried out on the Bionano Solve interface. Structural variations were highlighted by comparing optical maps results to *in silico* Col-0 TAIR10.1 reference genome labelling with DLE-1. A total of 797 SVs were identified during the analysis of L*er*-1 optical maps *versus* Col-0 TAIR10.1 reference (Table 1). When Bionano Solve tools detected one SV embedded in a second one, the largest SV was kept. This case was found on two Chr1 independent locations (INS:19 432 310..19 468 513 and DEL:24 688 666..24 736 849). A 1 kb size filter was applied on the Bionano SVs, that was equivalent to remove deletions and insertions with a Bionano quality score < 10 (defined as poor quality by the manufacturer) (Additional file 1: Table S15). Additionally, on Chr2, the INV SV (3 433 371..3 490 731) with no quality score was discarded. Thereby, 591 SVs representing 74.2% of total Bionano SVs were further considered in this analysis. INS and DEL types constituted the main part of the Bionano SVs (48.9% and 49.9% of the SVs respectively), the remaining 1.2% corresponding to translocations (TRA) and INV (Table 2). Median SVs size was 4 383 bp and SVs cumulated sizes represented 7.2 Mb of the genome. The TRA and INV types corresponded to nearly one third (2.0 Mb) of the structural variations cumulated size. In our study, the translocations were only detected using the Bionano assembly. The two L*er*-1 TRA were located on Chr2 (3 378 844..3 397 121; 3 484 209..3 844 839) (Additional file 2: Figure S6). The largest SV identified was a 1.1 Mb L*er*-1 INV located on Col-0 TAIR10.1 reference Chr4 (1 435 832..2 593 360) (Additional file 2: Figure S7). SVs were distributed preferentially along the chromosome arms and their detection was limited in centromeric regions due to decreased in labelling in these regions (Fig. 1).

### SVs comparison

SVs comparison was based on their absolute start- and end-positions on the Col-0 TAIR10.1 reference sequence. We considered structural variations were comparable in both technologies when their locations overlapped by at least 1 bp. To go further, SVs identified by ONT and Bionano technologies were assigned a two letters svID code with the first letter used for ONT SVs, the second for Bionano SVs, leading to common (svID UU and MU) and specific (svID UN and NU) locations (with “U” for “Unique SV”, “M” for “Multiple SVs” and “N” for “None SV”, Additional file 1: Tables S16 and S17).

SVs comparison metrics are presented in Table 3. A total of 563 common locations were identified representing 948 (80.1%) of ONT SVs and 563 (95.3%) of Bionano SVs. The cumulated sizes of these common SVs are 5.9 Mb and 6.9 Mb for ONT and Bionano detection respectively. ONT SVs tend to be smaller than Bionano SVs (Table 3, Additionnal File 1 Tables S16 and S17).

**Table 3.** Characteristics of Evry.L*er*-1 ONT and Bionano SVs identified in common and specific Col-0 TAIR10.1 locations.

| Technology | ONT |  | Bionano |  |
| --- | --- | --- | --- | --- |
|  | Common | Specific | Common | Specific |
| Locations | 563 | 236 | 563 | 28 |
| SVs > 1 kb (%) | 948 (80.1) | 236 (19.9) | 563 (95.3) | 28 (4.7) |
| Min Size | 1 003 | 1 003 | 1 034 | 1 017 |
| Max Size | 87 533 | 347 239 | 1 143 224 | 166 007 |
| Cumulated Size | 5.9 | 1.8 | 6.9 | 0.3 |
| Median Size | 3 759 | 2 656 | 4 456 | 1 374 |
| Average Size | 6 221 | 7 453 | 12 171 | 11 104 |

Note : Cumulated sizes are in Mb, and all other sizes in bp.

Among the 563 common regions, 410 (72.8% of the common regions) coincided with svID UU, *i.e.* one ONT structural variation corresponding to one SV Bionano. In most cases, the overlap of these SVs was at least 50.0% of the ONT SV size, and 405 (98.8% of the svID UU) SVs have conforming type (*i.e.* have the same type) (Additional file 1: Table S16). The remaining five svID UU (1.2%) were identified as deletions by ONT and insertions by Bionano technologies (svID UU_035, UU_038, UU_057, UU_073, UU_358; Additional file 1: Tables S16 and S17).

In the remaining 153 (27.2%) common locations, a total of 531 of the ONT SVs (56.8% of commons ONT SVs) related to the Bionano SVs (27.0% of commons Bionano SVs) were pinpointed (Table 4). These structural variations had a svID MU. The cumulative size of this SVs category is approximately 4 Mb for both technologies although the number of ONT variants is 3.5 times higher than in Bionano (531 vs 153). Nevertheless, Bionano median and average sizes are 2 and 4 fold larger respectively.

**Table 4.** Characteristics of the svID MU identified in ONT and Bionano SVs.

| Technology | ONT | Bionano |
| --- | --- | --- |
| Location | 153 | 153 |
| Number | 531 | 153 |
| Min Size | 1 010 | 1 253 |
| Max Size | 87 533 | 1 143 224 |
| Cumulated Size | 3.9 | 4.4 |
| Median Size | 4 236 | 9 523 |
| Average Size | 7 406 | 28 734 |

Note: svID MU corresponds to locations where Multiple ONT SVs overlap a Unique Bionano SV. Cumulated sizes are in Mb, and all other sizes in bp.

The largest ONT SV was a complex SV (svID MU_102; MU meaning that several ONT SVs match to one Bionano SV) consisting of four contiguous deletions located on Chr4. These four deletions coincided with one Bionano deletion (Additionnal file 1: Tables S16 and S17). The largest Bionano SVs (svID MU_097) was an inversion on Chr4 of 1 143 224 Mb overlapping 22 SV ONTs (corresponding to INS and DEL) (Additionnal file: 1 Table S17).

Specific locations were more abundant with the ONT technology (236 SVs - 19.9%) than with Bionano (28 SVs - 4.7%) leading to a cumulated size of 1.8 Mb and 0.3 Mb respectively, and with a median size twice larger (2 656 bp for ONT SVs *vs* 1 374 bp for Bionano SVs). The distribution of the specific ONT SVs onto the Col-0 TAIR10.1 chromosomes lead to a clear trend to locate on NOR and centromeres (Fig. 1). The largest specific ONT variant is located on Chr3 and corresponds to a DEL (svID UN_124; SV detected with ONT only, Additional file 1: Table S16). The largest specific Bionano SV is spotted on the Chr3 and corresponds to an INV type (svID NU_017; SV detected with Bionano only, Additional file 1: Table S17). A focus on the TRA revealed a 18.2 kb specific L*er*-1 Bionano SV (svID NU_007), close to the second TRA of 360 kb positioned around 3.6 Mb (MU_153). This TRA coincided with seven SV events (1 INV, 5 INS and 1 DEL) in the L*er*-1 SDN assembly (Additional file 1: Tables S16 and S17).

Using Araport11 annotation of the Col-0 reference (The Arabidopsis Information Resource - TAIR), a comparison using only ONT SVs is shown in Table 5. Since the Bionano events represent a large-scale observation, they were not taken into account in this analysis. A total of 893 (75.4%) out of 1184 ONT SVs overlaped TE features, of which 579 also overlaped genes. Only 291 (24.6%) SVs are located outside a TE feature, overlapping genes [125 (10.6%)] or not [166 (14.0%)] (Table 5). Focusing on ONT specific SVs (svID=UN), their overlap with the Col-0 reference annotation showed similar percentage compared to the common SVs.

**Table 5.**
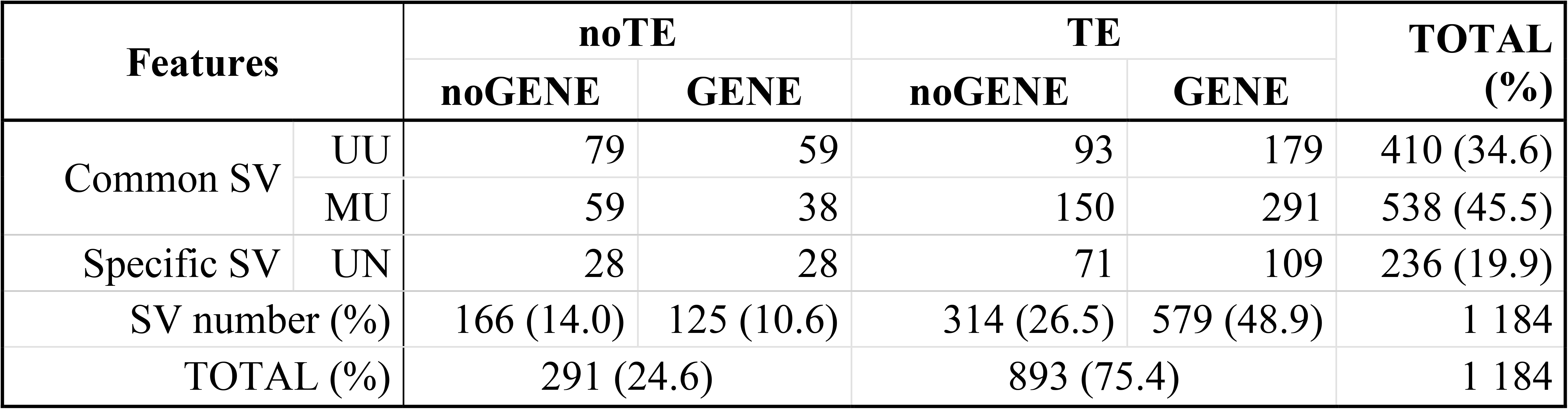
L*er*-1 ONT SVs (>1kb) overlapping Col-0 TAIR10.1 genes and TEs annotation features.

| Features |  | noTE |  | TE |  | TOTAL (%) |
| --- | --- | --- | --- | --- | --- | --- |
|  |  | noGENE | GENE | noGENE | GENE |  |
| Common SV | UU | 79 | 59 | 93 | 179 | 410 (34.6) |
|  | MU | 59 | 38 | 150 | 291 | 538 (45.5) |
| Specific SV | UN | 28 | 28 | 71 | 109 | 236 (19.9) |
| SV number (%) |  | 166 (14.0) | 125 (10.6) | 314 (26.5) | 579 (48.9) | 1 184 |
| TOTAL (%) |  | 291 (24.6) |  | 893 (75.4) |  | 1 184 |

To better characterize the genes affected by ONT SVs in common locations, a GO-terms overrepresentation test was performed with the PANTHER’s tool [61] available on TAIR website (https://www.arabidopsis.org/tools/go_term_enrichment.jsp). Among the 1 764 genes identified in common locations, 47.2% (832) genes were uniquely assigned to a GO term and used in PANTHER (Additional file 1: Tables S18 and S19). Overrepresentations in defense response and ADP-binding terms were detected (Additional file 1: Table S20), but no enrichment for GO-terms in genes in specific ONT locations was highlighted (Additional file 1: Tables S21-S23).

## Discussion

Herein, we compare the performance of Oxford Nanopore and Bionano Genomics technologies for structural variation detection. For this, we performed long read sequencing and optical mapping of two *A. thaliana* ecotypes, namely Columbia-0 (Col-0) and Landsberg *erecta* 1 (L*er*-1). Long read *de novo* assemblies were constructed using three different assemblers and optical maps were assembled with Bionano Solve tools. Structural variations detected using the Col-0 TAIR10.1 [58] and L*er* [44] genomic sequences as references, were described and compared to each other, to reveal the relative strengths of the two technologies in highlighting SVs.

### Assemblies based on ONT and Bionano data for SV analyses

To obtain the best assembly based on only long reads data we used three different assemblers. After comparison of assembly metrics, calculation time and collinearity against reference genomes, SDN provided the best assembly even if some collinearity breaks were observed, especially in centromeric regions. The metrics of Evry.Col-0 and Evry.L*er* 1 SDN assemblies were comparable to such assemblies in previous studies [24,44,45,62].

Continuous improvement in protocols and new developments in genome assembly strategies and algorithms resulted in higher and higher quality of genomic sequences used in subsequent analyses. Previously published Bionano *A. thaliana* optical map (KBS-Mac-74) genome [45] used a BspQI staining protocol for labelling, generating about 10 time more maps to cover the entire genome of KBS-Mac-74 than in our study (DLE-1 Bionano staining protocol), highlighting enhancement in Bionano’s protocol. In addition, no optical map was previously available for the Columbia (Col-0) and Landsberg *erecta* 1 (L*er*-1), making our map assemblies especially valuable for further studies.

Our high quality map allowed us to define centromeric and nucleolar organizer regions (NOR), despite lower molecules density and even if label concordance loss were observed between L*er*-1 maps compared to the Col-0 TAIR10.1 *in silico* reference maps. Moreover, fluctuations in ONT coverage density and accumulation of repetitive alignments in the same regions are reinforcing evidences of the approximate locations of the centromeres and NOR. However, we identified several missassemblies in the course of our SVs analyses between the ONT SDN L*er*-1 assembly and Col-0 TAIR10.1 reference, highlighting how difficult it can be to get a reliable assembly, and thus detecting SVs, in these complex regions.

### SV detection and comparison between the two technologies

Herein, we compared structural variations in Evry.L*er*-1 and the reference genome Col-0 TAIR10.1. We chose this reference because of its high quality and the richness of the associated studies [24,44,45].

The cumulated SVs sizes obtained for ONT and Bionano in our study are smaller than in previous studies [24,44]. Filtering on SVs size (SVs > 1kb *vs* no size filter) could explain this difference. In addition, the lack of duplications detection in ONT assembly could depend on MUMmer’s ability to detect this type of SV, reflecting the detection complexity of the duplication events, as mentioned in Goel *et al* (2019). In contrast, the absence of duplication detected by Bionano could be explained by polymorphic duplications between L*er*-1 maps and Col-0 TAIR10.1 reference, which would break the collinearity, as described in Jiao *et al.* (2020), and by the size of duplications (< 5kb, [62]) identified as the limit of Bionano detection. Analyzes by the two technologies revealed a predominance of insertion, deletion and inversion with larger median and average sizes for Bionano SVs. The distribution of these types of SV is homogeneous along the chromosomes arms. Even if most of the specific ONT SVs are located in the centromeric and pericentromeric regions, a decrease coverage of the SVs in these regions is probably due to technical problems such as assembly errors (for ONT SMARTdenovo). This diminution in SV coverage is also observed with Bionano technology, showing a lower density labeling in these complex regions. This contrasts with previous results identifying more SVs in regions where the recombination meiotic rate decreases [24]. The filtering of SV ONTs smaller than 1 kb could again be an explanation for this contradiction. On the other hand, Bionano Solve tools well identified translocation previously characterized on Chr2 and three inversions larger than 50 kb present on Chr3, Chr4 and Chr5 [24,35,44]. For example, compared to the Col-0 TAIR10.1 reference, the L*er*-1 maps support a 360 kb translocation of mitochondrial sequence in the Chr2 around the 3.6 Mb Col-0 TAIR10.1 position (svID MU_153). This observation is concordant with Stupar *et al.* (2001) that first described the mtDNA insertion in the Col-0 reference. In this Chr2 region (3.29 Mbp to 3.48 Mbp), Pucker *et al.* (2019) identified a second 300 kb highly divergent region between *A. thaliana* Nd-1 and Col-0 reference. In the same study, Pucker *et al.* also described the lack of the entire region between 3.29 Mbp and 3.48 Mbp in L*er* genome, corresponding to the specific translocation of 18.2 kb detected in L*er*-1 map (svID NU_007). Zooming in this Col-0 TAIR10.1 Chr2 region (3.2 Mb to 3.5 Mb) in the L*er*-1 SDN assembly, many small contigs are observed with a missing sequence of 110 kb. This observation explains absence of SV detection, confirming the great complexity of this region and the sequence divergence between L*er*-1 and Col-0 genome described by Pucker *et al* (2019). Even if the Col-0 reference sequence has been improved since 2000 [58], our assembly (Evry.Col-0) confirms its value to re-evaluate complex region assembly, and provide new high quality optical map data.

The number, type and location of SVs in the largest common ONT (svID MU_102) and Bionano (svID MU_097) SVs, as well as the Chr2 ONT SVs matching the second Bionano translocation (svID MU_153), reflect that the structural variations brought out by ONT were more numerous and smaller, which allows an identification at finer scale. In contrast, Bionano variants were larger and their sizes depend on restriction sites distribution.

To globally estimate consistency of the SV analyzes between ONT SDN or Bionano L*er*-1 assemblies against Col-0 TAIR10.1 reference, we compared the structural variations we identified to those of Zapata *et al.* (2016) (mapping and SV detection tools and parameters being the same). Although the local variations cannot be comparable due to genome sequence accuracy (complete genome *vs* whole genome sequencing) and the SV filtering differences (no size filter *vs* > 1 kb), the majority of events are shared by the both studies.

Comparing locations of the L*er*-1 ONT SVs with Araport11 annotations, we found that common and specific ONT SVs were preferentially linked to TE features and genes, as reported in Jiao *et al* (2020). Looking at the GO-term enrichment in genes overlapping common ONT SVs, an overrepresentation in defense response and ADP-binding terms corresponding to resistance genes was observed. This result is concordant with previous studies [13,24,44,63,64,65] in which an association between structural variations and the cluster organisation of resistance genes was described.

## General conclusion

Because analyses of SVs and their consequences heavily relies on the quality of their identification and the underlying assembly/mapping data, we aimed to compare the performance of ONT and Bionano biotechnologies for structural variation detection. Applying stringent filters on ONT assembly mapping approach and size filters on SVs, we have shown this methodology is an easy and efficient way to detect reliable SVs. Most of detected SVs were also identified with Bionano optical maps with high concordancy despite different characteristic (average, size, median). Nevertheless, long read sequencing technologies makes possible to detect SVs more accurately, while Bionano offers a broad overview of structural rearrangements. In addition, whole genome SVs analyses is currently mostly limited to model organisms. However, because both Oxford Nanopore long reads and Bionano Genomics maps assemblies do not require previous knowledge on genomic architecture or sequence of the studied taxa, this approach expands the field of suitable plant species or species complexes where in-depth SVs analyses can be performed.

Thereby, ONT appears to be especially suitable for SV studies in population or species complex, and Bionano more relevant for characterization of genome specificity and genome evolution, leading to an obvious complementarity of these two technologies in SVs analyses.

## Methods

### Plants

*Arabidopsis thaliana* Columbia-0 (accession number 186AV) and Landsberg *erecta*-1 (accession number 213AV) seeds were obtained from the Versailles Arabidopsis Stock Center, INRAE. They were sown directly in soil and transplanted after 10 days. Plantlets were grown under a 16h light/8 h night photoperiod in a growth chamber at 20°C for 4-5 weeks. Prior to harvest, the plants were dark-treated for 3 days.

### Oxford Nanopore Sequencing (MinION) HMW DNA extraction

High Molecular Weight (HMW) DNA extraction was performed using a modified salting-out protocol. A total of 5g of freshly harvested leaves was ground in liquid nitrogen with a mortar and pestle and transferred to 10ml of 50°C prewarmed extraction buffer in a 50ml tube containing 1.25% SDS, 100mM Tris-HCl, pH 8, 50mM EDTA, 0.01% w/v PVP40. Then 37.5μl of beta-mercaptoethanol (0.375% final) and 10μl RNAse A (Qiagen® 100mg/mL) were added. This solution was incubated for 30 min at 50°C, under agitation (10 sec at 300rpm every 10 min). After incubation, 20ml TE (10:1) were added, slowly homogenized then 10ml of KAc 5M. The tube was kept on ice for 5 minutes, then centrifuged at 4°C during 10 min at 5000g. The solution was transferred in two 15ml tubes and centrifuged again as previously. The supernatant was transferred in a 50ml tube contening 1 volume of Isopropanol, slowly inverted 10 times, then centrifuged at 4°C for 10min at 500g. Pellets were washed with 20ml ethanol 70% then centrifuged at 4°C for 5 min at 500g. Supernatant was removed and pellets were not completely dried before solubilization in 100μl of TE (10:1) prewarmed at 50°C. The DNA solution was then incubated at 50°C for 10 min. Field Inverted Gel Electrophoresis (Program 50-150 kb on *Pipin Pulse* from Sage Science) was used for DNA size estimation and DNA samples with molecule size above 50 kb were kept. Purity of DNA was evaluated by spectrophotometry (OD260/280 and OD260/230 ratio).

### Bionano Optical Maps ultra HMW DNA extraction

We performed the DNA extraction using the Base protocol n°30068 vD (Bionano Genomics) with minor adaptations. Three grammes of very young fresh leaves from each genotype were harvested from the dark-treated rosettes. The samples were placed on aluminium foil on ice then transferred to a 50ml tube surrounded by a screened cap allowing pouring without lost of samples (Bio-Rad) The tubes were kept on ice during the nuclear isolation. Samples were treated in fixing solution containing 2% formaldehyde under a fume hood then rinsed with fixing solution without formaldehyde. Fixed-leaves were transferred to a square Petri dish with 4ml of Plant Homogenization Buffer plus (HB+ is HB supplemented with 1mM spermine tetrahydrochloride, 1mM spermidine trihydrochloride, and 0.2% 2-mercaptoethanol). Entire leaves were chopped with a razor blade in 2×2mm pieces then transferred to a new tube on ice and 7.5ml HB+ is added. Using TissueRuptor (Qiagen) the 2×2mm pieces were blended for a total of four cycles (20 sec at maximum speed then resting 30 sec). Plant homogenates were filtered, first through a 100μm then to a 40μm cell strainer and volumes were adjusted to 45ml. Nuclei were centrifuged at 3840g at 4°C during 20 min, supernatants were discarded. Nuclei were gently re-suspended in residual buffer, 3ml of HB+ were added, then tubes were swirled on ice and the volumes were adjusted to 35ml. Homogenates were centrifuged at 60g at 4°C during 3 min using minimum deceleration. Solutions were very carefully transferred to a new tube in order to avoid carry-over of debris, and filtered again through a 40μm cell strainer.

Nuclei were centrifuged at 3840g at 4°C during 20 min, 3ml of HB+ were added and tubes were swirled on ice. Using Bionano Nuclei Purification by Density Gradient, nuclei homogenate were laid on the top of two solutions with different densities. After a 4500g centrifugation at 4°C during 40 min, the nuclei are at the interface of the two solutions. There are recovered with a wide-bore tip in about 1ml solution and transferred in a 15ml tube and adjusted to 14ml with HB+. Nuclei were centrifuged at 2500g at 4°C during 15 min. All the buffer were removed and nuclei were re-suspended in 60μl HB+.

The nuclei solution were adjusted to 43°C for 3 min and melted 2% agarose from CHEF Genomic DNA Plug Kits (Bio-Rad) was added to reach a 0.82% agarose plug concentration. Plugs were cooled on aluminum blocks refrigerated on ice. Purification of the plugs was performed with Bionano Lysis Buffer adjusted to pH 9 and supplemented with proteinase K and 0.4% 2-mercaptoethanol. Plugs were digested during 2h at 50°C in Thermomixer then solution were refreshed and incubated again overnight. Plugs were treated at RNAse for 1h at 37°C in remaining solution. Plugs were washed three times in Wash Buffer (Bionano Genomics) then four times in TE 10:1. DNA retrieval was performed as recommended by Bionano Genomics, as follow: plugs were melted at 70°C during 2 min then transferred immediately at 43°C and incubated 45 min at 43°C with 2μl Agarase (0.5 unit/μl). The melted plugs were recovered with wide-bore tips and dialyzed on a 0.1μm membrane disk (Millipore) floating on 10ml TE for 1h. DNA was quantified in triplicates with Qubit according to Bionano protocol. Two methods were used to estimate size of DNA molecules: *Pipin Pulse* and the Qcard Argus System (Opgen) which allows DNA combing on a lane and visualization of molecules after staining under fluorescent microscope. Samples with molecules above 150 kb were kept for labeling. Protocols were performed according to Bionano Genomics with 600ng of DNA for both Col-0 and L*er*-1 ecotypes. The direct label and stain (DLS) labeling consisted in a single enzymatic labelling reaction with DLE-1 enzyme following by DNA staining with a fluorescent marker. It was performed with 750ng DNA. Chip loading was performed as recommended by Bionano Genomics.

### ONT Sequencing (MinION) and assembly

ONT libraries were prepared according to the following protocol, using the Oxford Nanopore SQK-LSK109 kit. Genomic DNA or DNA previously fragmented to 50 kb with a Megaruptor (Diagenode S.A., Liege, Belgium) was first size-selected using a BluePippin (Sage Science, Beverly, MA, USA). The selected DNA fragments were end-repaired and 3’-adenylated with the NEBNext® Ultra™ II End Repair/dA-Tailing Module (New England Biolabs, Ipswich, MA, USA). The DNA was then purified with AMPure XP beads (Beckmann Coulter, Brea, CA, USA) and ligated with sequencing adapters provided by Oxford Nanopore Technologies (Oxford Nanopore Technologies Ltd, Oxford, UK) using Blunt/TA Ligase Master Mix (NEB). After purification with AMPure XP beads, the library was mixed with Running Buffer with Fuel Mix (ONT) and Library Loading Beads (ONT) and loaded on 4 MinION R9.4 SpotON Flow Cells per *Arabidopsis thaliana* ecotypes. Resulting FAST5 files were base-called using albacore (versions 2.1.10 and 2.3.1) and FASTA produced as described in Istace *et al* (2017). Canu version 1.5 (github commit ae9eecc), was used for initial read correction and trimming with the parameters minMemory=100G, corOutCoverage = 10000. The corrected sequences were merged in one final FASTA file per ecotype that were later used as assemblers input.

Assemblies were performed with the relevant genome size parameter set to, or coverage calculation based on, a 130 Mb genome size. Assemblers used with default parameters were Canu version 1.5 ([56], github commit 69b5f32), Rapid Assembler (RA, https://github.com/lbcb-sci/ra commit 07364a1) and SMARTdenovo version 1.0 (with the option –c 1 to run the consensus step) (https://github.com/ruanjue/smartdenovo commit 61cf13d). The MUMmer suite version 3.0 [59] was run with the parameters used in Zapata *et al*. 2016. To analyze the assemblies, they were aligned to the reference genome of *Arabidopsis thaliana* Columbia 0 (Col-0, TAIR10.1 GCF_000001735.4) and the sequence of *Arabidopsis thaliana* Landsberg *erecta* (L*er*, Genbank LUHQ00000000.1) using *nucmer* with the options -c 100 -b 500 -l 50 -g 100 -L 50. The alignments were filtered with *delta-filter* (options -1 -l 10000 -i 0.95) and visualized with the *mummer-plot* (options --fat --large --layout --png) or DNAnexus (github commit 78e3317). These MUMmer parameters [44] allowed conserving exact matches larger than 50bp and alignments longer than 10 kb with a minimal identity of 95%. To check assemblies’ completeness and fragmentation, they were compared to each other based on the metrics (Number of contigs, N50, cumulative genome sizes) and the genome alignments to the references generated with MUMmer viewed with the DNAnexus dot (https://dnanexus.github.io/dot/).

To evaluate the completness of our ONT data, mapping of the corrected ONT reads on the Col-0 TAIR10.1 reference were performed with Minimap2/2.15 aligner [57] with -a -x map-ont parameters. The Samtools/1.6 depth tool with –a option [66] gave us the alignement depth at each Col-0 TAIR10.1 reference positions.

### Bionano Optical Map assembly

As it can be beneficial for assembly steps, molecules sub-sampling was conducted when flowcells yielded more than 90 Gb and 600X of data. This adapted selection of molecules was made on each run with the Bionano RefAligner tool in command line (version 1.3.8041.8044 with –minlen 180 –randomize 1 –subset 1 nb_molec options) or with Bionano Access (version Solve3.3 with Filter Molecule Object utility) (Additional file 1: Tables S6 and S7).

Maps were then constructed with the tool *Generate de novo Assembly* of the Bionano Solve (version 3.3) using the options recommended by Bionano (With pre-assembly, Non haplotype without extend and split) and a 0.115 Gb genome size. The pre-assembly step calculates noise parameters that optimizes the quality of the assembly (less and larger maps). When a reference FASTA file is added, noise parameters are calculated in aligning the molecules to the reference.

Otherwise, the noise parameters are estimated thanks to a first rough assembly of the molecules. For Col-0 and L*er*-1 ecotypes, three maps were obtained, one without reference, one with the Col-0 reference and one with the L*er* reference (Additional file 1: Tables S8 and S9). In our study, the metrics of these assemblies are very similar. This stability reflects that noise parameters estimated either with references fasta sequences or our data, were comparable. This is a guaranty of quality of Bionano data and assemblies.

### ONT variation detection

Structural variations were obtained with MUMmer’s show-diff utility on the filtered alignments of SMARTdenovo assemblies against the references Col-0 and L*er*. One DIFF file per comparison were obtained. Six SV types (Gap, Duplication, Break, Jump, Inversion, Sequence) were described in the Additional file 2: Figure S4.

### Bionano variation detection

SVs detections were performed on the optical maps built with the public reference and our SMARTdenovo ONT assemblies using the tool Convert SMAP to VCF file. VCF files were recovered, describing all the structural variations between the optical maps and the considered reference. The variations were classified in 6 types: deletion, insertion, translocation and inversion. SVs detection stringency is intrinsic, based on the number of aligned molecules (at least nine by default) and the number of labels accross each variants breakpoint on the genome map (at least two by default) (Bionano tutorial : https://bionanogenomics.com/support-page/data-analysis-documentation/). The technology gave an interval with an uncertainty about breakpoint positions (CIPOS and CIEND in VCF files). In this study, these values were used to calculate the most extended positions for the Bionano SVs and avoid effect of label fluzz.

The low number of structural variations between Col-0 optical maps and the Col-0 TAIR10.1 reference (as L*er*-1 maps and L*er* reference) reflects the good collinearity between the map and the references. SVs gave us an indication on location of conflicts that could be due to mis-assemblies or intra-ecotype variations. Inter-ecotype detection allowed us to describe the variations between Col-0 and L*er*-1.

Quality and length characteristics were used to better describe and filter SVs. Bionano Solve associates a quality score to each INS and DEL based on sensitivity and the fraction of alternative calls in mix assemblies that were called in the alternative genome assembly [from no quality (.) or poor (0) to confident quality (20)]. We observed that this indicator follows the same trend as the SVs size (Additional file 1: Tables S11 and S15). Moreover, size range values where SVs abundances are the most different between both technologies are the extremes : the smallest (< 1 kb), where ONT technology detected much more SVs and the highest (> 5 kb) where Bionano technology detected proportionnally more SVs. So in our comparison analysis, to remove poor quality Bionano SVs, ONT sequencing errors and high sensitivity, a filter on query SV size (> 1 kp) was applied. Confidence scores for translocation and inversion breakpoints were computed as p-values, giving true confidence (in Mahalanobis distance) to positive calls. The recommended cutoffs are 0.1 and 0.01 for translocation and inversion breakpoints calls respectively and were used to eliminate uncertain inversion on Chr2.

### SV description

Custom-made R and Perl scripts were used to edit other tools outputs, describe ONT and Bionano SVs (types, size), locate SVs along the chromosomes and filter them. For ONT technology, SVs identified as assemblies discordances were quickly described and discarded before comparison. Those included sequences (SEQ), breaks (BRK) and jumps (JMP) ONT SV because they correspond to assembly or reference artefacts. Finally, size filters (more than 1 kb) were applied to take into accountONT high sequencing error rate, and low quality Bionano SVs. For Bionano SVs the largest absolute positions of the SV were conserved, taking into account the uncertainty around breakpoints due to the distance between two labels.

### SV comparison

Comparison of SV obtained with both ONT and Bionano technologies were based on the overlap of their absolute positions.

ONT SV and Bionano SVs files were used after conversion to BED format to identify overlapping regions with BEDtools (version 2.27.1, github commit cd82ed5, “bedtools intersect -wa -wb -a INPUT1.bed -b INPUT2.bed -loj > OUTPUT.bed”). Raw comparisons were then compared, compiled and formatted in one final output file using custom-made R scripts. For each SVs location, this file contained descriptors (SVs size, type, quality) for both technologies, information on the type of conflict and a 2 letter code. This code characterized the SVs location as follow : the first letter corresponds to the ONT SV characterization, the second to the Bionano SV. M (“Multiple”) means more than one SV, U (“Unique”) one SV, N (“None”) no SV. For example, the code “MU” means that this location arbored multiple ONT SV corresponding to a unique Bionano. The landscapes and SVs occurences visualization was performed with Circos/0.69.9 tool (perl/5.16.3) [67].

### SV and annotation

SVs overlapping a gene and/or TE were identified with the bedtools intersect by comparing their absolute positions to *A. thaliana* Col-0 annotations (11th july 2019 release, TAIR10_GFF3_genes_transposons.gff). Lists of genes impacted by SV for both technologies were extracted and a GO-term enrichment analysis performed using Fisher’s Exact test with a Bonferroni correction in PANTHER (released 20200407 with GO Ontology database DOI: 10.5281/zenodo.3873405 Released 2020-06-01, [61], http://go.pantherdb.org/). Significance was evaluated based on a P-value ≤ 10−5 and an FDR value ≤ 0.01 [67].

## List of abbreviations

If abbreviations are used in the text they should be defined in the text at first use, and a list of abbreviations can be provided.

bp: base pairs
BRK: Break
CGH: Comparative Genomic Hybridization
CNV: copy number variations
Col-0: *Arabidopsis thaliana* ecotypes Columbia-0
DEL: Deletion
DLE-1: Direct Label Enzyme – 1
DLS: Direct Label and Stain
DNA: Desoxyribo Nucleic Acid
DUP: Duplication
Gb: Gigabases
Hi-C: HIgh-throughput chromatin conformation Capture
Indels insertions/deletions
INS: Insertion
INV: Inversion
JMP: Jump
Kb: kilobases
L*er*-1: *Arabidopsis thaliana* ecotypes Landsberg *erecta* 1
LER: *Arabdopsis thaliana* L*er*-1 reference genome published by Zapata *et al*. 2016
NA: Not Available
NGS: Next Generation Sequence
ONT: Oxford Nanopore Technologies
PAV: presence/absence variations
RA: Rapid Assembler
SDN: SMARTdenovo
SEQ: Sequence
SNP: Single Nucleotid Polymorphism
SV: Structural Variation
TAIR10.1: last version of *Arabdopsis thaliana* Col-0 reference genome availbale at the The Arabidopsis Information Resource repository (TAIR)
TE: Transposable Element
TRA: Translocation

## Additional files description

**Additional_file_1.xlsx : Additional tables results**

Table S1. Metrics of the ONT run flowcells for *A. thaliana* Columbia (Col-0).

Note : All sizes are in base pairs. ALL_COL is all Col-0 trimmed merged data.

Table S2. Metrics of the ONT run flowcells for *A. thaliana* Landsberg *erecta* (L*er*-1). All sizes are in base pairs.

Note : All sizes are in base pairs. ALL_LER is all L*er*-1 trimmed merged data.

Table S3. Number of the Col-0 TAIR10.1 and L*er* references bases covered and uncovered by ONT reads.

Note : ONT reads are *A. thaliana* Col-0 and L*er*-1 corrected, trimmed and merged ONT sequences (respectively ALL_COL and ALL_LER).

Table S4. *A. thaliana* Col-0 Assembly Metrics for contigs only, obtained with SMARTdenovo, Canu and RA.

Note : All sizes are in base pairs. Assemblies are obtained with corrected, trimmed and merged Col-0 ONT sequences.

Table S5. *A. thaliana* L*er*-1 Assembly Metrics for contigs only, obtained SMARTdenovo, Canu and RA.

Note : All sizes are in base pairs. Assemblies are obtained with corrected, trimmed and merged L*er*-1 sequences.

Table S6. Metrics of the Bionano run chips for *A. thaliana* Columbia (Col-0) and Landsberg *erecta* (L*er*-1).

Note : All sizes are in base pairs. Results obtained with DLE-1 labelling.

Table S7. Metrics of the sampled Bionano run chips for *A. thaliana* Columbia (Col-0) and Landsberg *erecta* (L*er*-1).

Note : All sizes are in base pairs. Results obtained with DLE-1 labelling.

Table S8. Assembly Metrics of *A. thaliana* Columbia (Col-0) sampled molecules.

Note : The options used were “Pre-assembly”, “Non Haplotype” and “Without Extend and Split“.

Table S9. Assembly Metrics of *A. thaliana* Landsberg *erecta* (L*er*-1) sampled molecules.

Note : The options used were “Pre-assembly”, “Non Haplotype” and “Without Extend and Split”.

Table S10.Types of Col-0 ONT and Bionano SVs obtained against L*er* reference.

Table S11. Size repartition of Col-0 ONT and Bionano insertions, deletions, INVersions, translocations obtained against L*er* reference.

Table S12. Characteristics of Evry.Col-0 ONT and Bionano SVs, obtained after alignement against L*er* reference.

Table S13. Characteristics of compared ONT Col-0 SVs with query size > 1kb.

Table S14. Characteristics of compared Bionano Col-0 SVs with query size > 1kb.

Table S15. Size repartition of L*er*-1 ONT and Bionano insertions and deletions obtained against Col-0 TAIR10.1 reference.

Table S16. Characteristics of compared ONT L*er*-1 SVs with query size > 1kb.

Table S17. Characteristics of compared Bionano L*er*-1 SVs with query size > 1kb.

Table S18.Genes overlapping L*er*-1 SV in common locations (query size >1 kb).

Table S19. Gene annotation overlapping L*er*-1 SV in common locations (query size >1kb).

Note : PANTHER released 20200407 was used with GO Ontology database DOI: 10.5281/zenodo.3873405 Released 2020-06-01, [61], http://go.pantherdb.org/).

Table S20. PANTHER Overrepresentation results on Genes overlapping common L*er*-1 SVs (query size >1kb).

Note : The PANTHER version is decribed in Mi *et al*. 2019.

Table S21. Genes overlapping specific ONT L*er*-1 SVs (query size >1 kb).

Table S22. Gene annotation overlapping specific ONT L*er*-1 SVs (query size >1kb).

Note : PANTHER released 20200407 was used with GO Ontology database DOI: 10.5281/zenodo.3873405 Released 2020-06-01, [61], http://go.pantherdb.org/).

Table S23. PANTHER Overrepresentation results on Genes overlapping specific ONT L*er*-1 SVs (query size >1 kb).

Note : The PANTHER version is decribed in Mi *et al*. 2019

**Additional_file_2.pdf : Additional figures results :**

Figure S1A-C. Views of Col-0 contigs alignments on Col-0 TAIR10.1 reference (dotted end).

(A) Contigs obtained with SMARTdenovo, (B) with Canu and (C) with RA. Blue, green and orange dots and lines represent unique forward, unique reverse and repetitive alignments respectively.

Figure S2A-C. Views of L*er*-1 contigs alignments on L*er* reference (dotted end).

(A) Contigs obtained with SMARTdenovo, (B) with Canu and (C) with RA. Blue, green and orange dots and lines represent unique forward, unique reverse and repetitive alignments respectively.

Figure S3A-E. Bionano Access view of L*er*-1 cmaps aligned on Col-0 TAIR10.1 reference.

(A) to (E) are alignments on Col-0 TAIR10.1 Chr1 to Chr5. Maps are in green for the Col-0 TAIR10.1 reference and light blue for L*er*-1 genome with the molecules depth curve in blue. Consistant DLE-1 enzyme label between reference and L*er*-1 maps are represented with dark blue bars with grey links between the genomes maps. Inconsistant DLE-1 enzyme label are yellow bars on the two genomes maps.

Figure S4. Description of SVs detected by MUMmer show-diff and Bionano Access tools.

Insertion in the query are called GAP with a negative size by MUMmer show-diff, INS by Bionano Access. Deletion in the query are called GAP with a positive size by MUMmer show-diff, DEL by Bionano Access. Inversion in the query are called INV by MUMmer show-diff and Bionano Access. Duplication in the query are called DUP by MUMmer show-diff and by Bionano Access. Rearrangement of reference sequence in the query are called jump (JMP) by MUMmer show-diff and translocation (TRA) by Bionano Access. Inverted Duplication are not described by MUMmer show-diff and called INVDUP by Bionano Access. Reference sequence junction between two assemblies contigs alignment are called SEQ by MUMmer show-diff and are not described by Bionano Access. Query sequence junction between two reference chromosomes alignment are called break (BRK) by MUMmer show-diff and are not described by Bionano Access. « − » means no detection with the technology.

Figure S5A-E. Col-0 SVs (>1kb) occurences.

All comparisons were performed against the L*er* reference sequence per 100kb bins and black rectangles symbolize L*er* centromeric regions. Average mapping coverage for Col-0 ONT reads (red line called COV), average DLE-1 density labelling (green line called DLE), and ONT and Bionano occurrences (rea and green bars respectively) are represented for each L*er* chromosome in section A to E respectively for Chr1 to Chr5.

Figure S6. Bionano Solve zoom in the Chr2 L*er*-1 translocations against Col-0 TAIR10.1 reference.

Maps are in green for the Col-0 TAIR10.1 reference and light blue for L*er*-1 genome. Consistant DLE-1 enzyme label between reference and L*er*-1 maps are represented with dark blue bars with grey links between the genomes maps. Inconsistant DLE-1 enzyme label are yellow bars on the two genomes maps. The purple bar locate the translocation events on the Ler-1 map. The red box and lines highlight the zoom.

Figure S7. Bionano Solve capture of the L*er*-1 Chr4 extra-range Size Invertion against Col-0 TAIR10.1 reference.

Maps are in green for the Col-0 TAIR10.1 reference and light blue for L*er*-1 genome. Consistant DLE-1 enzyme label between reference and L*er*-1 maps are represented with dark blue bars with grey links between the genomes maps. Inconsistant DLE-1 enzyme label are yellow bars on the two genomes maps. The red box and lines highlight the zoom.

## Declarations

### Ethics approval and consent to participate

Not applicable.

### Consent for publication

Not applicable.

### Availability of data and materials

The ONT reads files and the Bionano molecules files have been submitted to the European Nucleotide Archive (http://www.ebi.ac.uk) and are publicly available with the accession numbers ERP128342 and ERZ1959921 respectively. Assemblies and optical maps of the Col-0 and L*er*-1 genomes are publicly available in separate ENA studies under the accession number PRJEB44316.

### Authors’ contributions

The project was conceived by VB, MCLP and PFR. Plant cultures and the HMW DNA extraction for Oxford Nanopore and Bionano Technologies were carried out by ED and GM, data acquisition by ED, GM, CC, CB, BI and equipment provided by PW, MCLP, PFR, VB. Data analysis were performed by AC, BI and CB for assemblies, optical maps and SV detection and AC and RG for SV comparisons. PFR, VB contributed to data interpretation with AC and RG. The manuscript was written by AC, RG, PFR, VB with inputs from PW and MCLP. All authors read and approved the final manuscript.

## Acknowledgements

The authors would like to thank the Versailles *Arabidopsis* Stock Center, INRAE providing *Arabidopsis thaliana* ecotypes and the Institut of Plant Sciences Paris Saclay (IPS2) where the *Arabidopsis* cultures were carried out.

We also greatly thank Damien Hinsinger for proofreading and advice on the manuscript and the INRAE PepiAnnot Group (https://pepi-ibis.inra.fr/annotation-genomes) for helpful discussions.

## Footnotes

Not applicable.

## References

1. Saxena RK, Edwards D, Varshney RK. Structural variations in plant genomes. Brief Funct Genomics. 2014 Jul;13(4):296–307.

2. Escaramís G, Docampo E, Rabionet R. A decade of structural variants: description, history and methods to detect structural variation. Brief Funct Genomics. 2015 Sep 1;14(5):305–14.

3. Zhang X, Chen X, Liang P, Tang H. Cataloging Plant Genome Structural Variations. Current Issues in Molecular Biology. 2018;181–94.

4. Ho SS, Urban AE, Mills RE. Structural variation in the sequencing era. Nature Reviews Genetics. 2020 Mar;21(3):171–89.

5. Wendel JF, Jackson SA, Meyers BC, Wing RA. Evolution of plant genome architecture. Genome Biol. 2016 Dec;17(1):1–14.

6. Gabur I, Chawla HS, Snowdon RJ, Parkin IAP. Connecting genome structural variation with complex traits in crop plants. Theor Appl Genet. 2019 Mar 1;132(3):733–50.

7. Schiessl S-V, Katche E, Ihien E, Chawla HS, Mason AS. The role of genomic structural variation in the genetic improvement of polyploid crops. The Crop Journal. 2019 Apr;7(2):127–40.

8. Voichek Y, Weigel D. Identifying genetic variants underlying phenotypic variation in plants without complete genomes. Nat Genet. 2020 May;52(5):534–40.

9. Muñoz-Amatriaín M, Eichten SR, Wicker T, Richmond TA, Mascher M, Steuernagel B, et al. Distribution, functional impact, and origin mechanisms of copy number variation in the barley genome. Genome Biol. 2013 Jun;14(6):R58.

10. Dolatabadian A, Patel DA, Edwards D, Batley J. Copy number variation and disease resistance in plants. Theor Appl Genet. 2017 Dec 1;130(12):2479–90.

11. Fuentes RR, Chebotarov D, Duitama J, Smith S, De la Hoz JF, Mohiyuddin M, et al. Structural variants in 3000 rice genomes. Genome Res. 2019;29(5):870–80.

12. Tao Y, Zhao X, Mace E, Henry R, Jordan D. Exploring and Exploiting Pan-genomics for Crop Improvement. Molecular Plant. 2019 Feb 4;12(2):156–69.

13. Wei H, Liu J, Guo Q, Pan L, Chai S, Cheng Y, et al. Genomic Organization and Comparative Phylogenic Analysis of NBS-LRR Resistance Gene Family in Solanum pimpinellifolium and Arabidopsis thaliana. Evol Bioinform Online. 2020;16:1176934320911055.

14. Prunier J, Caron S, MacKay J. CNVs into the wild: screening the genomes of conifer trees (Picea spp.) reveals fewer gene copy number variations in hybrids and links to adaptation. BMC Genomics. 2017 Dec;18(1):97.

15. Prunier J, Giguère I, Ryan N, Guy R, Soolanayakanahally R, Isabel N, et al. Gene copy number variations involved in balsam poplar (Populus balsamifera L.) adaptive variations. Mol Ecol. 2019 Mar;28(6):1476–90.

16. Wang Y, Xiong G, Hu J, Jiang L, Yu H, Xu J, et al. Copy number variation at the GL7 locus contributes to grain size diversity in rice. Nature Genetics. 2015 Aug;47(8):944–8.

17. Gong C, Du Q, Xie J, Quan M, Chen B, Zhang D. Dissection of Insertion–Deletion Variants within Differentially Expressed Genes Involved in Wood Formation in Populus. Front Plant Sci [Internet]. 2018 [cited 2019 Aug 20];8. Available from: https://www.frontiersin.org/articles/10.3389/fpls.2017.02199/full?report=reader

18. Gao L, Gonda I, Sun H, Ma Q, Bao K, Tieman DM, et al. The tomato pan-genome uncovers new genes and a rare allele regulating fruit flavor. Nature Genetics. 2019 Jun;51(6):1044.

19. Tranchant-Dubreuil C, Rouard M, Sabot F. Plant Pangenome: Impacts On Phenotypes And Evolution. Annual Plant Reviews [Internet]. 2019 May [cited 2021 Feb 11]; Available from: https://hal.archives-ouvertes.fr/hal-02053647

20. Khan AW, Garg V, Roorkiwal M, Golicz AA, Edwards D, Varshney RK. Super-Pangenome by Integrating the Wild Side of a Species for Accelerated Crop Improvement. Trends in Plant Science. 2020 Feb 1;25(2):148–58.

21. Li R, Li Y, Zheng H, Luo R, Zhu H, Li Q, et al. Building the sequence map of the human pan-genome. Nature Biotechnology. 2010 Jan;28(1):57–63.

22. Sherman RM, Forman J, Antonescu V, Puiu D, Daya M, Rafaels N, et al. Assembly of a pan-genome from deep sequencing of 910 humans of African descent. Nature Genetics. 2019 Jan;51(1):30–5.

23. Duan Z, Qiao Y, Lu J, Lu H, Zhang W, Yan F, et al. HUPAN: a pan-genome analysis pipeline for human genomes. Genome Biology. 2019 Jul 31;20(1):149.

24. Jiao W-B, Schneeberger K. Chromosome-level assemblies of multiple Arabidopsis genomes reveal hotspots of rearrangements with altered evolutionary dynamics. Nature Communications. 2020 Feb 20;11(1):989.

25. Song J-M, Guan Z, Hu J, Guo C, Yang Z, Wang S, et al. Eight high-quality genomes reveal pan-genome architecture and ecotype differentiation of Brassica napus. Nat Plants. 2020 Jan;6(1):34–45.

26. Springer NM, Ying K, Fu Y, Ji T, Yeh C-T, Jia Y, et al. Maize Inbreds Exhibit High Levels of Copy Number Variation (CNV) and Presence/Absence Variation (PAV) in Genome Content. PLOS Genetics. 2009 Nov 20;5(11):e1000734.

27. Swanson-Wagner RA, Eichten SR, Kumari S, Tiffin P, Stein JC, Ware D, et al. Pervasive gene content variation and copy number variation in maize and its undomesticated progenitor. Genome Res. 2010 Jan 12;20(12):1689–99.

28. Hwang JE, Kim S-H, Jung IJ, Han SM, Ahn J-W, Kwon S-J, et al. Comparative genomic hybridization analysis of rice dwarf mutants induced by gamma irradiation. Genet Mol Res. 2016 Dec 23;15(4).

29. Gimode D, Odeny DA, de Villiers EP, Wanyonyi S, Dida MM, Mneney EE, et al. Identification of SNP and SSR Markers in Finger Millet Using Next Generation Sequencing Technologies. PLoS ONE. 2016;11(7):e0159437.

30. Lyons E, Freeling M, Kustu S, Inwood W. Using Genomic Sequencing for Classical Genetics in E. coli K12. PLoS One [Internet]. 2011 Feb 25 [cited 2020 May 15];6(2). Available from: https://www.ncbi.nlm.nih.gov/pmc/articles/PMC3045373/

31. Brenchley R, Spannagl M, Pfeifer M, Barker GLA, D’Amore R, Allen AM, et al. Analysis of the bread wheat genome using whole genome shotgun sequencing. Nature. 2012 Nov 29;491(7426):705–10.

32. Xing L, Zhang D, Song X, Weng K, Shen Y, Li Y, et al. Genome-Wide Sequence Variation Identification and Floral-Associated Trait Comparisons Based on the Re-sequencing of the ‘Nagafu No. 2’ and ‘Qinguan’ Varieties of Apple (Malus domestica Borkh.). Front Plant Sci [Internet]. 2016 [cited 2020 May 11];7. Available from: https://www.frontiersin.org/articles/10.3389/fpls.2016.00908/full

33. Maldonado dos Santos JV, Valliyodan B, Joshi T, Khan SM, Liu Y, Wang J, et al. Evaluation of genetic variation among Brazilian soybean cultivars through genome resequencing. BMC Genomics. 2016 Feb 13;17(1):110.

34. The 1001 Genomes Consortium, Alonso-Blanco C, Andrade J, Becker C, Bemm F, Bergelson J, et al. 1,135 Genomes Reveal the Global Pattern of Polymorphism in Arabidopsis thaliana. Cell. 2016 Jul 14;166(2):481–91.

35. Pucker B, Holtgräwe D, Rosleff Sörensen T, Stracke R, Viehöver P, Weisshaar B. A De Novo Genome Sequence Assembly of the Arabidopsis thaliana Accession Niederzenz-1 Displays Presence/Absence Variation and Strong Synteny. Vandepoele K, editor. PLoS ONE. 2016 Oct 6;11(10):e0164321.

36. Pinosio S, Giacomello S, Faivre-Rampant P, Taylor G, Jorge V, Le Paslier MC, et al. Characterization of the Poplar Pan-Genome by Genome-Wide Identification of Structural Variation. Mol Biol Evol. 2016 Oct;33(10):2706–19.

37. Redmond SN, Sharma A, Sharakhov I, Tu Z, Sharakhova M, Neafsey DE. Linked-read sequencing identifies abundant microinversions and introgression in the arboviral vector Aedes aegypti. BMC Biol. 2020 Dec;18(1):26.

38. Rhoads A, Au KF. PacBio Sequencing and Its Applications. Genomics Proteomics Bioinformatics. 2015 Oct;13(5):278–89.

39. Lu H, Giordano F, Ning Z. Oxford Nanopore MinION Sequencing and Genome Assembly. Genomics Proteomics Bioinformatics. 2016 Oct;14(5):265–79.

40. Mahmoud M, Gobet N, Cruz-Dávalos DI, Mounier N, Dessimoz C, Sedlazeck FJ. Structural variant calling: the long and the short of it. Genome Biology. 2019 Nov 20;20(1):246.

41. Huddleston J, Chaisson MJP, Steinberg KM, Warren W, Hoekzema K, Gordon D, et al. Discovery and genotyping of structural variation from long-read haploid genome sequence data. Genome Res. 2017 Jan 5;27(5):677–85.

42. De Coster W, D’Hert S, Schultz DT, Cruts M, Van Broeckhoven C, Berger B. NanoPack: visualizing and processing long-read sequencing data. Bioinformatics. 2018 Aug 1;34(15):2666–9.

43. Chaisson MJP, Sanders AD, Zhao X, Malhotra A, Porubsky D, Rausch T, et al. Multi-platform discovery of haplotype-resolved structural variation in human genomes. Nature Communications. 2019 Apr 16;10(1):1784.

44. Zapata L, Ding J, Willing E-M, Hartwig B, Bezdan D, Jiao W-B, et al. Chromosome-level assembly of *Arabidopsis thaliana* L *er* reveals the extent of translocation and inversion polymorphisms. Proceedings of the National Academy of Sciences. 2016 Jul 12;113(28):E4052–60.

45. Michael TP, Jupe F, Bemm F, Motley ST, Sandoval JP, Lanz C, et al. High contiguity Arabidopsis thaliana genome assembly with a single nanopore flow cell. Nature Communications. 2018 Feb 7;9(1):541.

46. Belser C, Istace B, Denis E, Dubarry M, Baurens F-C, Falentin C, et al. Chromosome-scale assemblies of plant genomes using nanopore long reads and optical maps. Nature Plants. 2018 Nov;4(11):879–87.

47. Sun S, Zhou Y, Chen J, Shi J, Zhao H, Zhao H, et al. Extensive intraspecific gene order and gene structural variations between Mo17 and other maize genomes. Nat Genet. 2018 Sep;50(9):1289–95.

48. Lam ET, Hastie A, Lin C, Ehrlich D, Das SK, Austin MD, et al. Genome mapping on nanochannel arrays for structural variation analysis and sequence assembly. Nat Biotechnol. 2012 Aug;30(8):771–6.

49. Cao H, Hastie AR, Cao D, Lam ET, Sun Y, Huang H, et al. Rapid detection of structural variation in a human genome using nanochannel-based genome mapping technology. GigaSci. 2014 Dec;3(1):34.

50. Levy-Sakin M, Pastor S, Mostovoy Y, Li L, Leung AKY, McCaffrey J, et al. Genome maps across 26 human populations reveal population-specific patterns of structural variation. Nature Communications. 2019 Mar 4;10(1):1025.

51. Leung AK-Y, Liu MC-J, Li L, Lai YY-Y, Chu C, Kwok P-Y, et al. OMMA enables population-scale analysis of complex genomic features and phylogenomic relationships from nanochannel-based optical maps. Gigascience [Internet]. 2019 Jul 1 [cited 2019 Sep 24];8(7). Available from: https://academic.oup.com/gigascience/article/8/7/giz079/5530323

52. Soto DC, Shew C, Mastoras M, Schmidt JM, Sahasrabudhe R, Kaya G, et al. Identification of Structural Variation in Chimpanzees Using Optical Mapping and Nanopore Sequencing. Genes (Basel). 2020 Mar 4;11(3).

53. Yuan Y, Milec Z, Bayer PE, Vrána J, Doležel J, Edwards D, et al. Large-Scale Structural Variation Detection in Subterranean Clover Subtypes Using Optical Mapping. Front Plant Sci. 2018 Jul 17;9:971.

54. Dixon JR, Xu J, Dileep V, Zhan Y, Song F, Le VT, et al. Integrative detection and analysis of structural variation in cancer genomes. Nat Genet. 2018 Oct;50(10):1388–98.

55. Long E, Evans C, Chaston J, Udall JA. Genomic Structural Variations Within Five Continental Populations of Drosophila melanogaster. G3 (Bethesda). 2018 Aug 15;8(10):3247–53.

56. Koren S, Walenz BP, Berlin K, Miller JR, Bergman NH, Phillippy AM. Canu: scalable and accurate long-read assembly via adaptive *k* -mer weighting and repeat separation. Genome Research. 2017 May;27(5):722–36.

57. Li H. Minimap2: pairwise alignment for nucleotide sequences. Bioinformatics. 2018 Sep 15;34(18):3094–100.

58. The Arabidopsis Genome Initiative. Analysis of the genome sequence of the flowering plant Arabidopsis thaliana. Nature. 2000 Dec 1;408(6814):796–815.

59. Kurtz S, Phillippy A, Delcher AL, Smoot M, Shumway M, Antonescu C, et al. Versatile and open software for comparing large genomes. Genome Biology. 2004;9.

60. Jain M, Tyson J, Loose M, Ip C, Eccles D, O’Grady J, et al. MinION Analysis and Reference Consortium: Phase 2 data release and analysis of R9.0 chemistry. F1000Research. 2017 May 31;6:760.

61. Mi H, Muruganujan A, Ebert D, Huang X, Thomas PD. PANTHER version 14: more genomes, a new PANTHER GO-slim and improvements in enrichment analysis tools. Nucleic Acids Res. 2019 Jan 8;47(D1):D419–26.

62. Goel M, Sun H, Jiao W-B, Schneeberger K. SyRI: finding genomic rearrangements and local sequence differences from whole-genome assemblies. Genome Biol. 2019 Dec;20(1):277.

63. Meyers BC, Kozik A, Griego A, Kuang H, Michelmore RW. Genome-wide analysis of NBS-LRR-encoding genes in Arabidopsis. Plant Cell. 2003 Apr;15(4):809–34.

64. Song Y, Ling N, Ma J, Wang J, Zhu C, Raza W, et al. Grafting Resulted in a Distinct Proteomic Profile of Watermelon Root Exudates Relative to the Un-Grafted Watermelon and the Rootstock Plant. J Plant Growth Regul. 2016 Sep 1;35(3):778–91.

65. Staal J, Kaliff M, Bohman S, Dixelius C. Transgressive segregation reveals two Arabidopsis TIR-NB-LRR resistance genes effective against Leptosphaeria maculans, causal agent of blackleg disease. Plant J. 2006 Apr;46(2):218–30.

66. Li H, Handsaker B, Wysoker A, Fennell T, Ruan J, Homer N, et al. The Sequence Alignment/Map format and SAMtools. Bioinformatics. 2009 Aug 15;25(16):2078–9.

67. Krzywinski M, Schein J, Birol I, Connors J, Gascoyne R, Horsman D, et al. Circos: An information aesthetic for comparative genomics. Genome Research. 2009 Sep 1;19(9):1639–45.

68. Lin M, Fang J, Hu C, Qi X, Sun S, Chen J, et al. Genome-wide DNA polymorphisms in four Actinidia arguta genotypes based on whole-genome re-sequencing. PLOS ONE. 2020 Apr 10;15(4):e0219884.

69. Toda N, Rustenholz C, Baud A, Le Paslier M-C, Amselem J, Merdinoglu D, et al. NLGenomeSweeper: A Tool for Genome-Wide NBS-LRR Resistance Gene Identification. Genes (Basel) [Internet]. 2020 Mar 20 [cited 2021 Apr 12];11(3). Available from: https://www.ncbi.nlm.nih.gov/pmc/articles/PMC7141099/

