## Additional file 2 for "Oxford Nanopore and Bionano Genomics technologies evaluation for plant structural variation detection"

Figure S1. Views of Col-0 contigs alignments against Col-0 TAIR10.1 reference (dotted end).

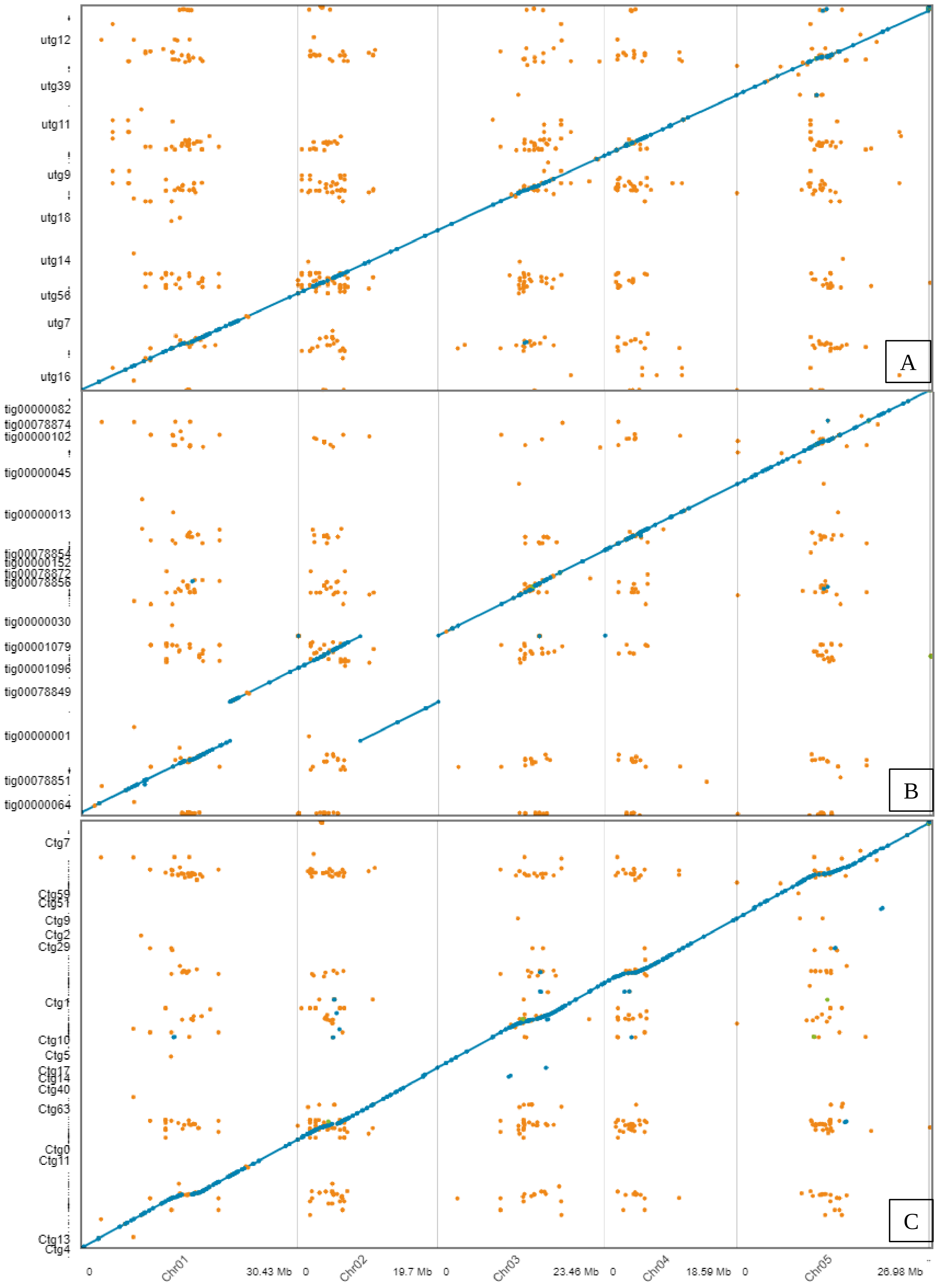

Figure S2. Views of L*er*-1 contigs alignments on L*er* reference (dotted end).

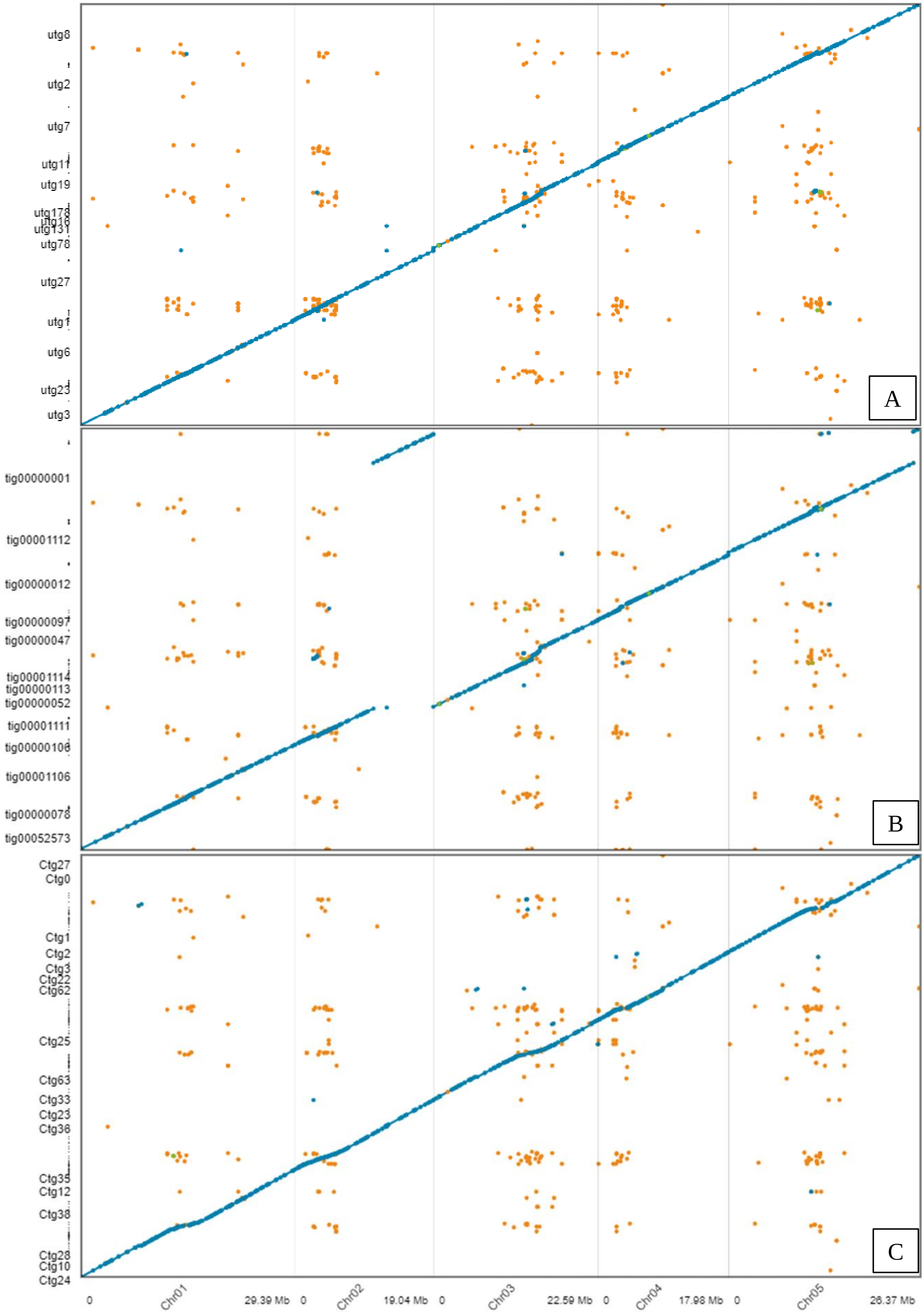

Figure S3. Bionano Access view of L*er*-1 maps aligned on Col-0 TAIR10.1 reference.

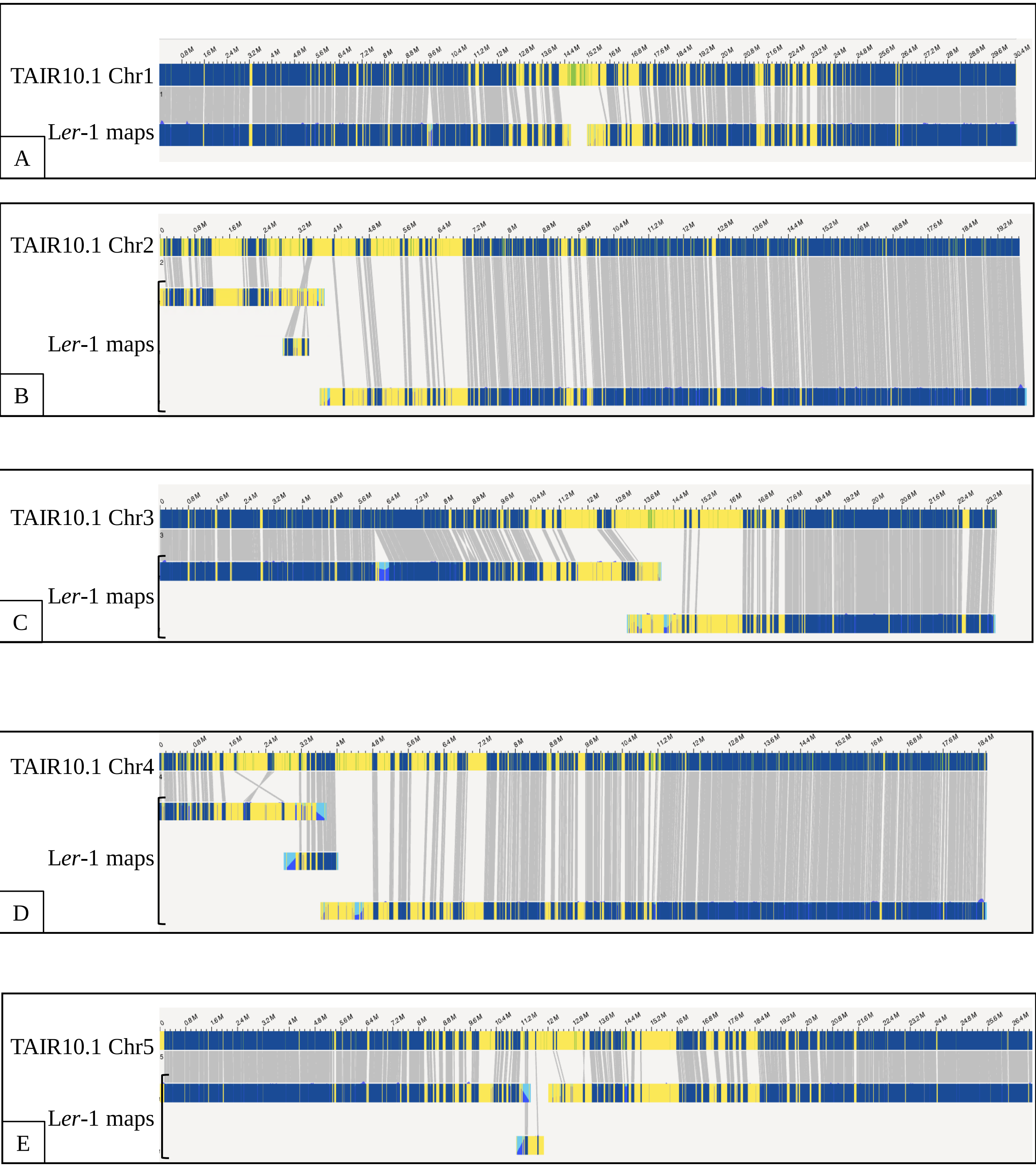

Figure S4. Description of SVs detected by MUMmer show-diff and Bionano Access tools.

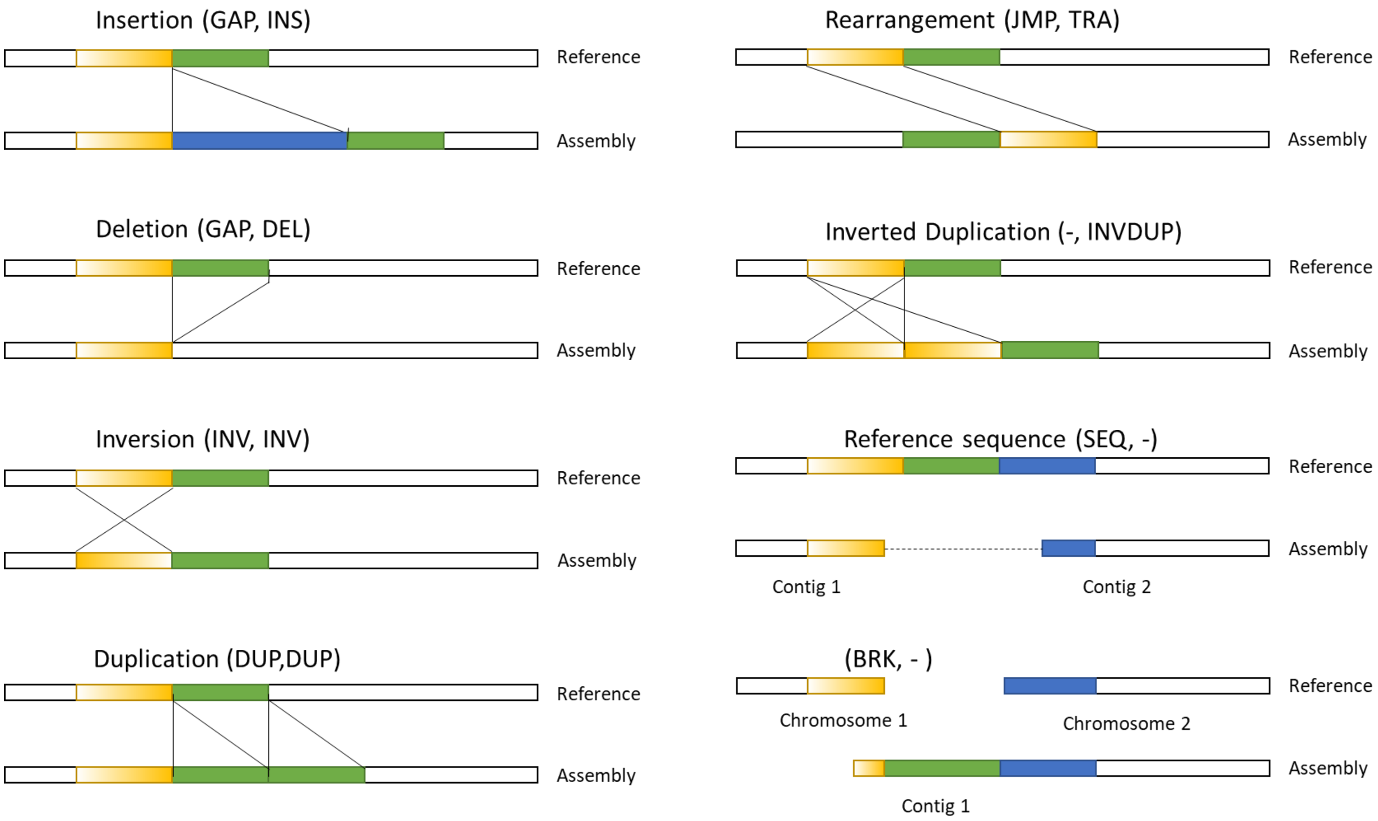

Figure S5. Col-0 SVs (>1kb) occurrences and landscape of L*er* reference chromosomes.

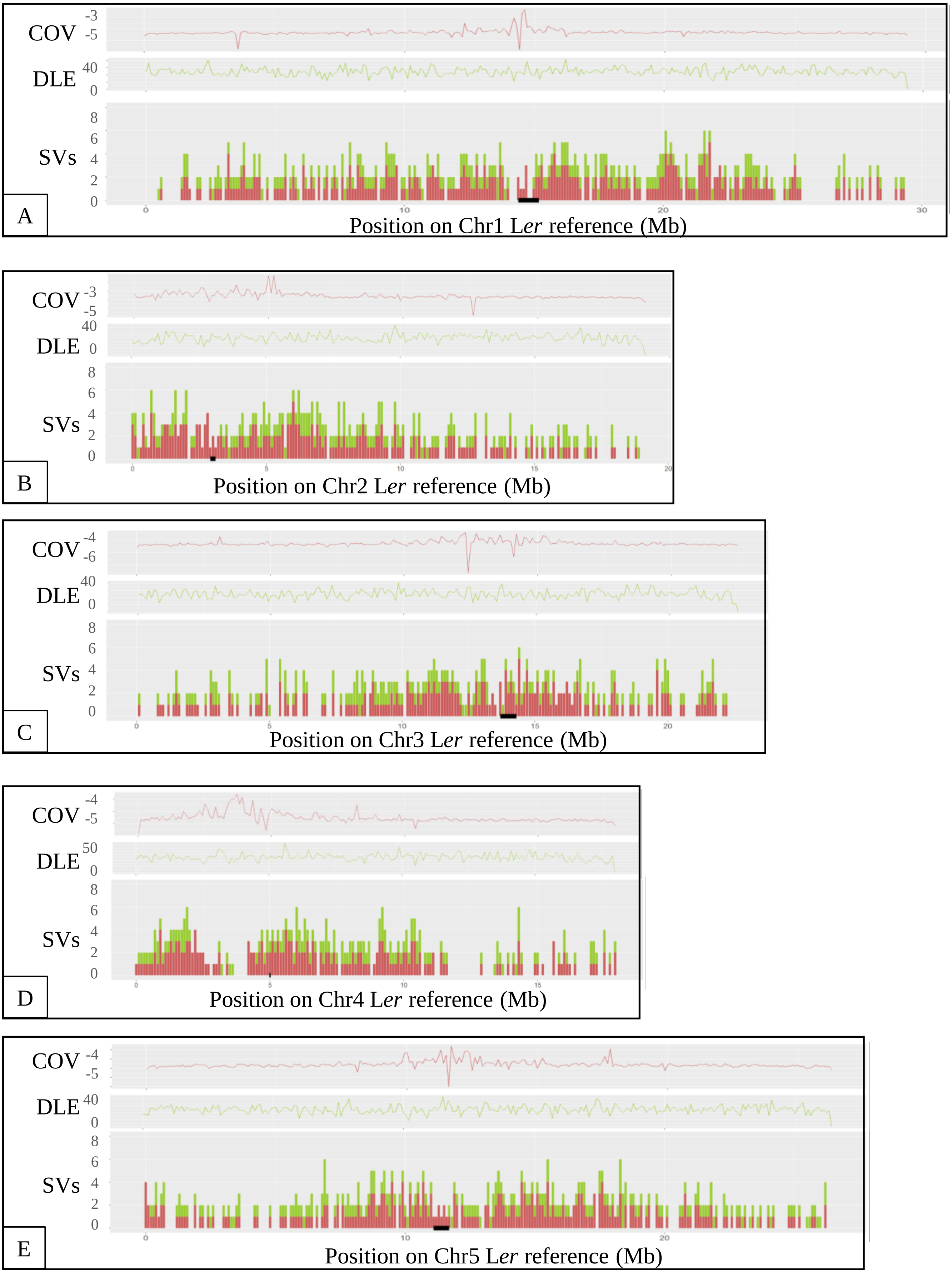

Figure S6. Bionano Solve zoom in the Chr2 L*er*-1 translocations against Col-0 TAIR10.1 reference.

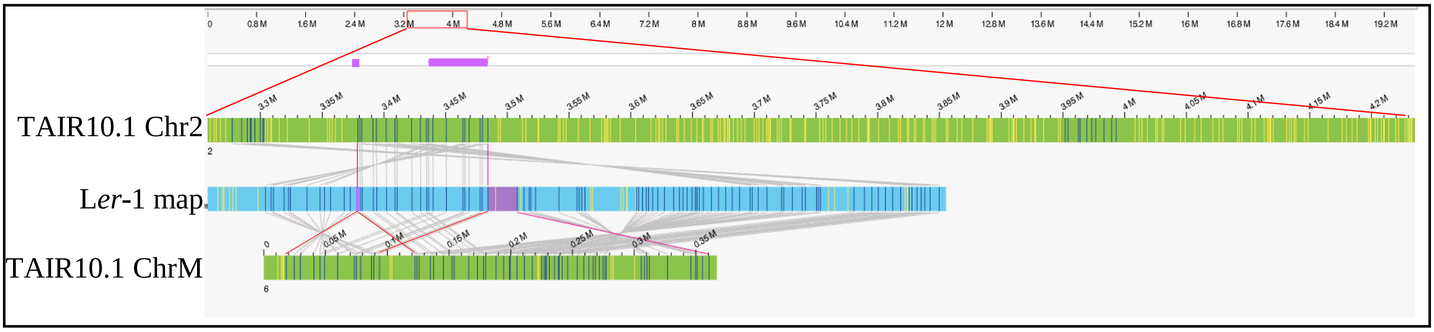


Figure S7. Bionano Solve capture of the L*er*-1 Chr4 extra-range size invertion against Col-0 TAIR10.1 reference.

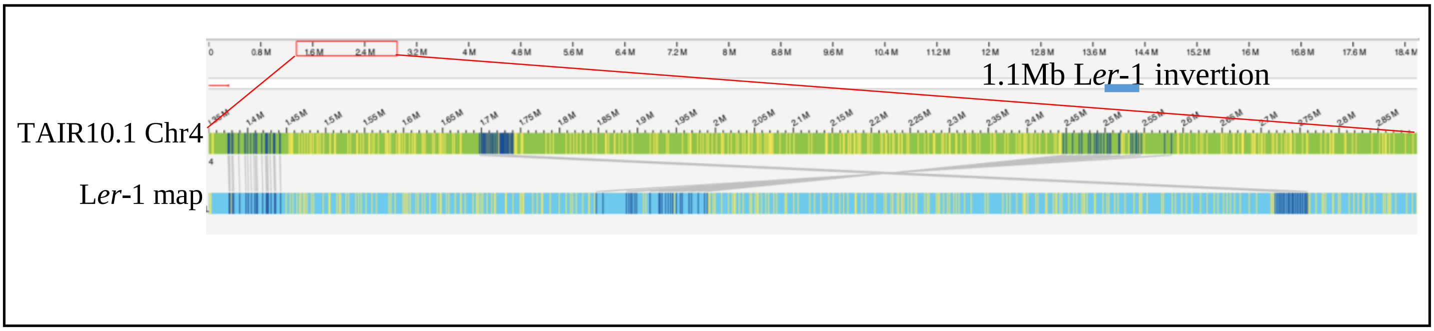
